# Towards an analytical framework for investigating the neurophysiological influence of social context in speech perception

**DOI:** 10.64898/2026.11.05.764590

**Authors:** Emily Y.J. Ip, Aoife Igoe, Benjamin R. Cowan, Giovanni M. Di Liberto

## Abstract

Social information plays an important role in everyday speech communication, yet existing neurophysiological approaches provide limited means of isolating its contribution during naturalistic speech perception. Here, we present an analytical framework that combines the large language model (LLM) Mistral-Instruct, fine-tuned on conversational language, and temporal response functions (TRFs) to investigate the influence of social context on word-level speech processing. The framework disentangles local context (LC) from speaker-specific social context (SC) when generating LLM-derived lexical expectation measures, enabling such predictions to be generated under either matching or mismatching speaker contexts for the same speech material. This provides a direct way of isolating the contribution of social information to lexical predictions. We first show that speaker-related social context carries information about speech register and conversational role beyond that available from previous local context alone by interrogating Mistral-Instruct. We then demonstrate that lexical expectations generated from socially congruent contexts produce small but reliable improvements in EEG prediction relative to expectations generated from incongruent social contexts. These effects were observed despite identical acoustic input and neural recordings, indicating that the influence of social context on lexical expectations explains dissociable variance in neural responses to speech. Together, these findings provide a proof of concept for using LLM-derived lexical expectations to isolate and measure the influence of social context on speech processing. More broadly, the proposed framework offers a flexible approach for generating and testing hypotheses about how different forms of contextual knowledge shape neural responses during naturalistic language comprehension.

## 1. Introduction

Speech is a fundamental means of human communication, enabling the exchange of information, the expression of thoughts, and the coordination of social behavior. Recent advances in continuous speech paradigms have provided important insights into how the brain derives multiple levels of linguistic representation and meaning from a single acoustic speech trace (Di Liberto et al., 2023; Doelling et al., 2014; Lesenfants et al., 2019; O’Sullivan et al., 2015; Vanthornhout et al., 2018). One phenomenon that has emerged from this work is cortical tracking, the alignment of neural activity with properties of ongoing speech, which has become a key framework for characterizing the neural mechanisms underlying speech perception and comprehension (Di Liberto et al., 2022; Obleser & Kayser, 2019). However, a large body of cortical tracking research has focused only on how the brain processes single streams of speech, such as audiobooks and stories (Broderick et al., 2018; Broderick et al., 2021; Di Liberto et al., 2015; Ding & Simon, 2012; Dou et al., 2024; Gillis et al., 2021; Heilbron et al., 2022; Piazza et al., 2025). In contrast, everyday communication is inherently social: the processing and understanding of speech depends not only on linguistic input, but also on a speaker’s communicative intentions, shared knowledge in *common ground*, a speakers’ beliefs and biases, the relationship between interlocutors, and social-cognition skills (Brennan et al., 2010; Brennan & Metzing, 2004). These social factors can profoundly shape what we say (underlying meaning) and how we say it (word form), yet much remains unknown about their influence on the neural mechanisms captured by cortical tracking.

Here, we propose an analytic approach for measuring how social information influences the neural processing of speech at the lexical level. To validate the methodology, we designed an electroencephalography (EEG) experiment involving a speech listening task focusing on one particular type of social information: speech register. Speakers routinely adapt their language to the characteristics and needs of their interlocutor (Clark, 1996; Clark & Murphy, 1982; Giles & Ogay, 2007). The resulting adapted speech registers in the adult-adult and adult-child conversations capture a well-established form of communicative accommodation. To that end, in this study we specifically consider two conditions with strong word-level differences: adult-directed speech (ADS) and child-directed speech (CDS) (Rowe & Snow, 2020; Soderstrom, 2007). Intuitively, on top of other particularities, for example at the supra-segmental level (e.g., prosody), CDS is expected to involve a simpler lexicon, with less variation and the use of particular selection of word forms (e.g., less over-complicated technical terms) (Fernald, 1989; Ferreira, 2019). Our experiment uses dialogue recordings from popular podcasts including dialogues between adult dyads or adult-child dyads, where the EEG participant takes the perspective of an *unremarked* third-person listener. Our numerical analysis also takes into account that differences in speech register can also be accompanied by differences in disfluency profiles (Ip et al., 2025), exploring the robustness of our method to the presence of more or less frequent disfluency, informing us on the applicability of this methodology in more realistic conversational scenarios.

First, we explored if typical cortical tracking measurements are affected by speech register in this specific dataset using the temporal response function (TRF) method (Crosse et al., 2016a). This analysis has one caveat: ADS and CDS can differ across multiple dimensions, so changes in a given cortical tracking measure (e.g., sound envelope tracking) may be underpinned both by acoustic differences and top-down effects of higher-order differences, such as lexical level differences (Synigal et al., 2026). To solve that issue, we propose a second approach to isolate the influence of social information on word-level processing. Specifically, we examine whether information about the speaker context is incorporated into listeners’ next-word predictions, and whether these socially informed predictions are reflected in neural responses to ongoing speech.

We propose isolating the influence of social information on word-level processing by combining TRFs and large language models (LLMs). We take a predictive processing view, positing that prior context influences the neural processing and representation of upcoming words (Clark, 2013; DeLong et al., 2005; Federmeier, 2007; Friston, 2005; Kuperberg & and Jaeger, 2016). Previous studies have operationalized contextual influences on lexical processing using measures including word dissimilarity derived from distributional embeddings (Broderick et al., 2018) and next-word expectations derived from LLMs (Heilbron et al., 2022). However, these approaches have predominantly characterized expectations based on prior proximal context, or local context (LC), with some exceptions (Yeshurun et al., 2017). Here, we additionally incorporate speaker-specific context, which we refer to more broadly as social context (SC), reflecting the type of conversation, and specifically the ADS/CDS contrast. By manipulating the context supplied to the LLM, the proposed approach isolates the unique contribution of SC and tests whether socially-informed lexical expectations are reflected in the neural response to continuous speech.

## 2. Methods

### 2.1 Experimental design

Twenty healthy native English speakers (10 female; age range 20-39 years, mean = 25.9 years, SD = 4.9 years) were recruited in Dublin for the experiment. Participants reported no history of neurological disorders or hearing impairment, provided written informed consent, and received financial compensation for their participation. Ethical approval was obtained from the School of Psychology ethics committee at Trinity College Dublin, and all procedures conformed to the Declaration of Helsinki.

The listening materials consisted of 42 naturalistic podcast dialogues (total duration ≈ 1 h) drawn from publicly available online sources. These included episodes from *Brains On!* and *Forever Ago*, *WIRED 5 Levels of Difficulty* series on YouTube, and additional interviews featuring the *Forever Ago* hosts, available online^1^. The recordings covered general-interest topics such as science and mathematics and were identical across participants.

Sixteen recordings featured conversations between two adults (AA), whereas twenty-six involved adult-child interactions (AC). The same five adult speakers appeared across both conditions, speaking with different children and adult interlocutors. Because child-directed and child-produced speech tends to occur at a slower speaking rate than adult-adult conversation, more AC recordings were included to achieve a comparable number of words across conditions (Ferreira, 2019; Snow, 1972). To that end, the selected speech registers captured a range of speaking styles for testing the robustness of our findings (e.g., the impact of disfluencies on the EEG encoding of speech) to variations in acoustic and lexical properties of a speech input.

Participants were presented with the forty-two speech stimuli (referred hereafter as trials) in a random order, separated by a short break. To encourage attention, participants were asked to rate how interesting they found the speech segment on a scale from 1-5 (where 1 = not at all interesting, 3 = neutral, 5 = very interesting) after each trial. Additionally, for a random subset of twenty percent of the trials, the participants were asked a behavioral multiple-choice question to select the set of keywords that best described the trial they just heard. This was also to keep the participants engaged and on task throughout the experiment.

The EEG dataset analyzed in the present study was collected as part of a previously reported speech listening experiment (Ip et al., 2025). Key aspects of the data acquisition procedure are summarized here. EEG was recorded using a 64-channel BioSemi ActiveTwo system at a sampling rate of 512 Hz. Auditory stimuli were delivered through Sennheiser HD 280 Pro headphones (44.1 kHz) using PsychoPy version 2022.2.5 (Peirce et al., 2019). Participants completed the experiment in a dimly lit room at Trinity College Dublin while fixating a centrally presented cross and minimizing movement.

### 2.2 EEG data preprocessing

Neural data were analyzed offline using MATLAB software (MathWorks). EEG signals were digitally filtered using Butterworth zero-phase filters with order 2, implemented with the function *filtfilt*. The primary analyses were carried out on EEG data band-pass filtered between 0.5 and 8 Hz, as that range has been shown to be particularly sensitive to the cortical tracking of speech (Di Liberto et al., 2015; Doelling et al., 2014; O’Sullivan et al., 2015; Vanthornhout et al., 2018). Low-frequencies from 0.5 Hz to 1 Hz were included as they were previously shown to be important for measuring the neural encoding of lexical predictions with EEG (Broderick et al., 2018; Carta et al., 2024). EEG signals were down sampled to 64 Hz, reducing computational and storage costs. EEG channels with a variance exceeding three times that of the surrounding ones were replaced by a spherical spline interpolation (Delorme & Makeig, 2004; Perrin et al., 1989). Signals were then re-referenced to the average of the two mastoid channels.

### 2.3 Stimulus feature extraction

A previous analysis of this dataset indicated a robust cortical tracking of speech envelope, envelope derivative, and word onset in response to the podcast recordings (Ip et al., 2025), in line with previous EEG and MEG work (Brodbeck et al., 2018; Di Liberto et al., 2015; Ding & Simon, 2012; Gillis et al., 2021; Lalor & Foxe, 2010; Lalor et al., 2009; Power et al., 2012). Our previous analysis also showed that LLM-estimations of next-word surprisal and entropy are also robustly encoded in the EEG signal. The present study goes beyond that by testing if these next-word expectations are built only using local proximal context (LC) or a combination of LC and social context (SC).

Word surprisal and entropy are sparse time-series, consisting primarily of zero values, where non-zero values correspond to word onsets derived using a forced aligner (see (Ip et al., 2025) for further details). Word surprisal and entropy time-series are built by scaling the non-zero values corresponding to word onsets with the LLM-derived word surprisal and entropy values respectively. These two expectation measures are complementary aspects of the same prediction. Word entropy indexes the uncertainty over possible upcoming words prior to their occurrence, capturing the brain’s state before the next-word is received and the distribution of the prediction. In contrast, word surprisal reflects how much an upcoming word deviates from those expectations, indexing prediction error.

Word expectations were built using the Open-Source LLM Mistral-Instruct-7B-v.01 model. Such measures are typically extracted by prompting the model with the words from the previous proximal context to get a set of probabilities for the upcoming word, which are transformed into surprisal and entropy accordingly. To incorporate SC into the lexical expectation model, we first adapted the model input to represent dialogue structure and then manipulated the SC information supplied alongside LC (**Figure 1**). Two additions provided the dialogue framing: 1) speaker-tags and 2) a system prompt identifying the discourse as dialogue. A final addition of 3) SC was then manipulated independently to generate congruent and incongruent lexical expectations.

**Figure 1.**
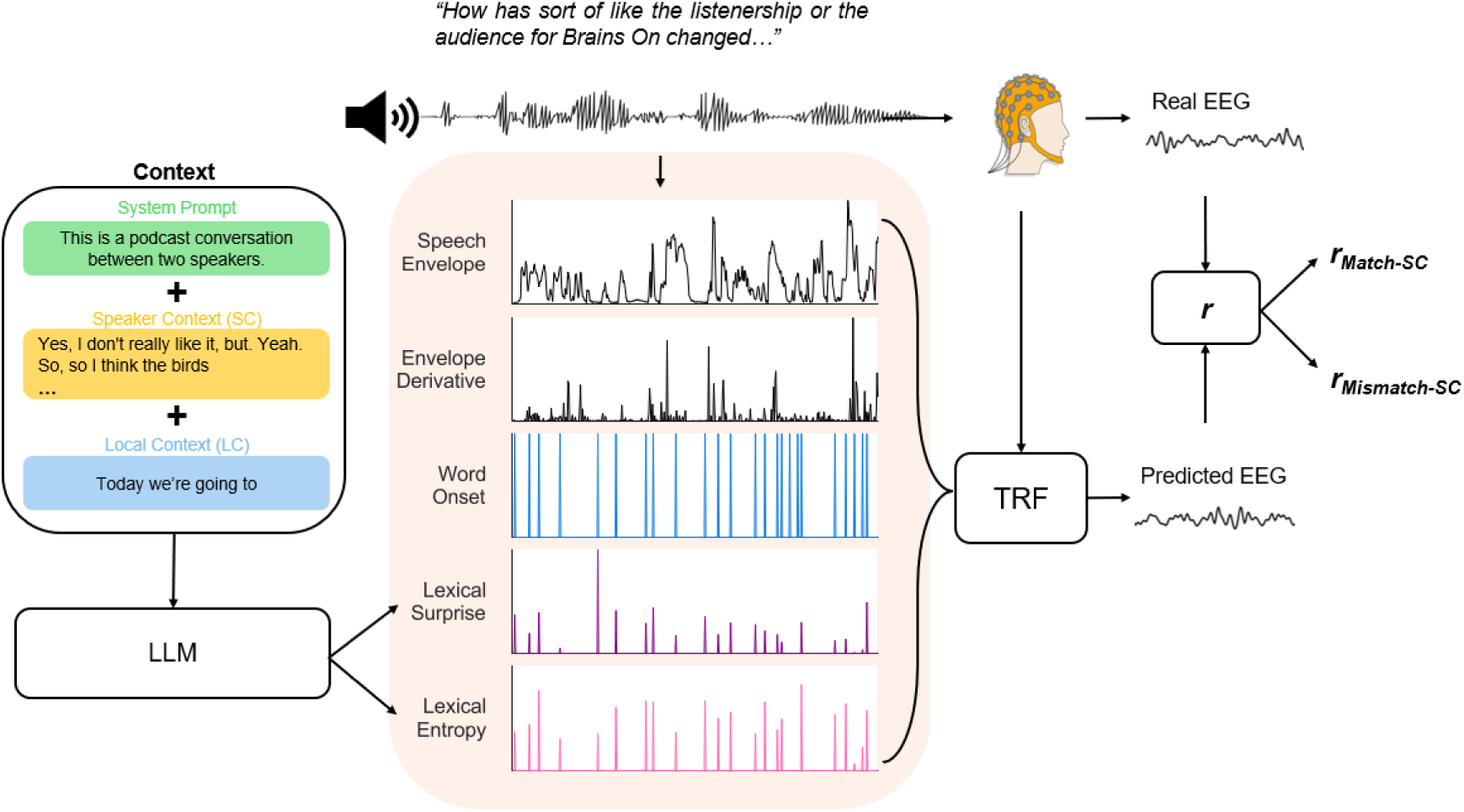
Experimental design and analysis framework. Neural signals were recorded as participants listened to natural, spontaneous speech from podcast dialogue recordings. The dialogues were represented with acoustic (speech envelope, envelope derivative, word onset) and lexical (surprisal and entropy) speech features, which were linearly regressed with the low-frequency EEG signals (0.5-8 Hz) to fit the temporal response function models.

1. *Speaker tags* were added to the transcripts (i.e., “Speaker 1”, “Speaker 2”) before each turn to provide Mistral with information about speaker identity and turn structure. These tags remained agnostic to specific speaker characteristics (e.g., “host/guest”, “adult/child”), avoiding the introduction of additional assumptions about speaker roles during prediction.
2. An explicit *system prompt* identifying the input as a podcast conversation between two speakers was provided to condition the model on a dialogue context: “*This is a podcast conversation between two speakers*”. Whereas the speaker tags identified who was speaking and when turns changed, the system prompt provided an explicit discourse-level framing through which the model could interpret this structure as dialogue. Prior work on prompt conditions indicate that system-level instructions and conversational formatting influence a language model’s next-token predictions. Therefore, the prompt was intended to steer Mistral’s predictions towards patterns characteristic of conversational interaction (Deng et al., 2023; Swamy et al., 2023), lowering the overall surprisal towards items like “yeah/so/like” that are common in turn-taking behaviors.
3. Inclusion of social information in the prompt. To that end, we subdivided the prompt into the two distinct segments SC and LC. LC includes words immediately preceding the current word, keeping predictions on-topic of the current conversation and aware of the turns being taken in the dialogue. SC could be defined in many ways and future studies may explore other approaches. Here, SC consisted of a random selection of utterances from different podcast recordings from the same register (note that podcast recordings differ in the topic of conversation, limiting the value of SC to speech register and conversational role). We grouped text according to four registers: adult-directed speech from the podcast host (ADS_host_) and guest (ADS_guest_), child-directed speech (CDS_host_), and child speech (CS_guest_; note that adult-child conversations always included a child guest and adult host). Based on the categorical divisions outlined in **Figure 2**, we built SC by concatenating a random selection of phrases from the same speech register (SC_match_) and compared with a baseline constructed using a different speech register (SC_mismatch_).

One central consideration was the balance between the lengths of LC and SC. Previous work deriving lexical expectations from LLMs has typically utilized contexts (i.e., LC) including all previous words possible in a given speech segment (Caucheteux et al., 2022; Goldstein et al., 2022; Heilbron et al., 2022; Schrimpf et al., 2021). These lexical expectations were also generated on the current dataset, where the segments have an average word count of 247 words (Ip et al., 2025). However, long segments of prior context is likely to contain not only local linguistic information but also information about speaker register and conversational role, reducing the unique contribution of an additional SC. The rationale here is to maximize that prediction gain, so that the contribution of social information to the prediction and the corresponding neural correlates can be isolated. To that end, we explored varied proportions of LC and SC sizes to identify an LC-SC ratio that is optimal for isolating the influence of social information on next-word expectations. One caveat is that LC should be sufficiently small to be (mostly) oblivious to social information. An LC that is too small, however, would make the prediction meaningless, thus raising the need for identifying a sweet-spot between prediction quality and social information in LC.

The explored LC and SC sizes were between 1 to 64, scaling logarithmically. These sizes were selected as they are consistent with the range of context lengths over which Mistral makes meaningful inferences (Mistral AI, 2026) and because such window sizes more closely resemble the plausible limits of human short-term memory of prior linguistic context, in contrast to the current ‘superhuman’ memories of LLMs (Oh & Linzen, 2026). Additionally, empirical evidence from neural encoding models with fMRI demonstrates a ceiling of effective prediction modeling with LLMs to brain activity at a window of 32 tokens (Tikochinski et al., 2025), with performance declining for larger windows. The ranges of LC and SC are also consistent with the types of utterances made during dyadic speech. At the lower end, the turn-taking nature of conversation often leads to utterances in short segments (e.g., 3-5 words), as seen in responses expressing agreement (“yeah” or “no”) or affective reactions (“huh,” “woah,” “really?”). At the same time, natural dialogue may also include longer utterances of speech in which interlocutors elaborate more extensively on their opinions to establish common ground.

### 2.4 Temporal response function

To characterize the relationship between speech features and neural activity, we applied a Temporal Response Function (TRF) modelling approach (Ding & Simon, 2012; Lalor et al., 2009). The TRF is a system-identification framework that uses regularized linear regression to estimate how speech features of interest are encoded in EEG brain activity over time (Crosse et al., 2016a). In a *forward* model, speech features are used to predict neural activity. The resulting TRF, a linear filter, represents a set of time-lagged weights that describes how the brain responds to properties of the speech signal.

The TRF provides two interpretable measures of the cortical tracking of speech. First, the model weights describe the temporal evolution of the neural response to a given speech feature. Because neural responses occur in reaction to a given stimulus with delays rather than instantaneously, the model estimates a series of time-lagged weights that capture how stimulus information influences EEG activity across subsequent latencies. These weights can be examined to identify the timing and scalp distribution of speech-evoked neural processes.

Second, model performance is quantified through prediction accuracy. A leave-one-out cross-validation was used to evaluate model generalizability (Crosse et al., 2016a; Crosse et al., 2021). For each fold, the TRF was trained on all trials except one and then used to reconstruct the EEG signal from the held-out trial. Prediction accuracy was calculated as the Pearson correlation between the predicted and recorded EEG signals at each electrode. Higher prediction correlations indicate that a greater proportion of EEG variance is explained by the speech feature(s) included in the model, providing an index of the strength of neural encoding. For each participant, prediction correlations were averaged across all EEG electrodes to obtain subject-level values for use in all statistical tests. We report the EEG prediction correlations calculated using the **GT-TC** (*ground-truth time-constrained*) metric, combining the approaches proposed in (Chalehchaleh et al., 2025; Di Liberto et al., 2023), and tailored to word level processing, as presented by Chalehchaleh et al., 2025.

In the present study, we performed a within-condition comparison within each speech register using multivariate TRF models fit on both acoustic features and lexical expectation features. Importantly, this comparison did not involve any manipulation of the stimulus or neural data. Instead, the same speech was represented using lexical expectation features generated under either congruent or incongruent speaker contexts. Consequently, any differences in encoding performance reflect the extent to which neural activity tracks contextually congruent lexical expectations.

### 2.5 LLM analyses

Having established the SC+LC procedure, we next assessed what (social) information an LLM could infer from each context type. To do so, for each SC size, a set of 10,000 SCs were randomly sampled. LLM Mistral was prompted with each of those SC segments at a time, interrogating the model explicitly with forced-choice questions on the register and role of the speaker. This procedure was also repeated for LC, providing a valuable comparison (**Figure S2**).

Transcripts were annotated for their speaker role (host/guest) and speaker register (child/adult). The model “`mistral-small-latest`” was selected for this analysis, which provides a publicly accessible API. Mistral-small was prompted as follows:

> *“Analyze the following utterance, making inferences about the socially-relevant information in its context, and answer each question by selecting ONLY from the options provided in parentheses.*

*Questions:*

1. *What is the speaker register? (adult / child)*
2. *Is this a dialogue or monologue? (monologue / dialogue)*
3. *What is the role of the current speaker? (host / guest)”*

The text results were parsed via custom code to collect the answers, producing the results in **Figures 3 and S2.**

### 2.6 Statistical analyses

Statistical analyses were performed in MATLAB. Mistral classification performance (Register and Role) was assessed against chance level (50%) using one-tailed binomial tests, with p-values corrected for multiple comparisons using the Benjamini-Hochberg false discovery rate (FDR) procedure (α = 0.05) (Benjamini & Hochberg, 1995). Differences in classification accuracy between local and social context were assessed using two-sided two-proportion z-tests, with the same FDR correction applied across context sizes. EEG prediction correlations derived from the multivariate TRF models, averaged across scalp conditions, were compared between experimental conditions (ADS vs CDS/CS) using two-sided paired Wilcoxon signed-rank tests separate for the envelope and word onset features. To assess whether encoding was significantly above zero, the prediction correlations for each feature and condition were tested against zero using one-sided Wilcoxon signed-rank tests. The effect of experimental condition on TRF feature weights at channel CZ was assessed at each time lag using paired Wilcoxon signed-rank tests, with p-values FDR-corrected across time lags (α = 0.05).

To measure effects across social context sizes and congruency conditions (congruent vs incongruent SCs), prediction correlations were tested using two-way repeated measures ANOVAs. Where sphericity assumptions were violated (*p* < 0.05), Greenhouse-Geisser corrections were applied to the significance tests. Significant omnibus effects were followed by pairwise Wilcoxon signed-rank tests with FDR correction (α = 0.05). To assess effects across electrodes and time (-200 to 600 ms) on the prediction correlations, one-sided paired cluster-based permutation tests (Maris & Oostenveld, 2007) were performed using the *permutest* MATLAB toolbox (Gerber, 2026), with significance determined from 10,000 permutations. Spatially adjacent electrodes exceeding an uncorrected paired-sample threshold (α = 0.05) in the hypothesized direction (Match > Mismatch) were grouped into clusters. The summed cluster-level test statistic (T-sum) was compared against a null distribution generated from 10,000 permutations of the condition labels within subjects to identify electrode clusters that were responsible for significant prediction gains in the congruent models over the incongruent models. This procedure was repeated independently for each SC size to assess the robustness of the prediction gains across SC sizes.

### 2.7 Data availability

The TRF analysis was conducted using the publicly available and open source mTRF-toolbox version 2.6 (Crosse et al., 2016b; Crosse et al., 2021), which can be downloaded from https://github.com/mickcrosse/mTRF-Toolbox. Stimulus features and EEG data were stored according to the continuous-event neural data structure CND (Di Liberto et al., 2024). Analyses were conducted using scripts shared via the GitHub channel^2^ of the Cognition and Natural Sensory Processing (CNSP) and Natural Sensory Processing open science initiative^3^. The analysis code for the standard forward TRF analysis can be found at the GitHub channel under the script titled *CNSP_example1_forwardTRF_multipleConditions.m.* Custom scripts were used for data visualization and statistical analyses. The dataset analyzed in this study is publicly available via the open science repository OSF (osf.io/rv6fw) and in the list of publicly available datasets of the CNSP-initiative.

## 3. Results

A multivariate TRF model was fit between the EEG signal and the envelope and word onset features for each condition (ADS and CDS/CS). No statistically significant differences were detected when comparing EEG prediction correlations (Wilcoxon paired signed-rank test (\**p* <0.05): envelope: *p* = 0.167; word onset: *p* = 0.156). TRF weights at Cz differed significantly between AA and AC in a cluster spanning 140-172 ms for the envelope feature (FDR-corrected Wilcoxon signed-rank test, *p* = 0.029), but not for the word onset feature (all FDR-corrected *p* > 0.05) (**Figure S1**). This lack of difference in traditional metrics for this specific contrast further motivates the development of a more tailored analytic procedure. Our rationale was that the responses may be similar across conditions on average, but the expectations at the individual word level would be different across conditions. Here, we compare SC match and mismatch to measure that contrast.

Next, we investigated the effectiveness of the new SC+LC. The first step was to ensure that the LC used in the LLM procedure captures the least amount of social information as possible. We interrogated the LLM Mistral to determine the largest possible LC size that is also (mostly) insensitive to speaker register and role. *Speaker role* (host vs. guest) classification for LC was only above chance level for size 32 (FDR-corrected, one-tailed binomial test vs chance (*p =* 0.5), across all context sizes and both context types: *p* < 0.001), indicating that any size below 32 is acceptable. *Speech register* (ADS vs. CDS/CS) was instead detectable above chance across the full range of LC sizes tested 9 FDR-corrected, one-tailed binomial test vs chance (*p =* 0.5), across all context sizes and both context types: *p* < 0.001) (**Figure S2B**). Nonetheless, an LC of size 4 had a lower sensitivity than the SC equivalent (LC 62.7% vs. SC 70.5%; two-proportion z-test, *z* = −3.408, FDR-corrected *p* < 0.001). On this basis, we selected an LC size of 4 words for all subsequent analyses, to best compromise between having a low next-word surprisal (**Figure S2A**). We also include supplementary results for LC of size 8, which would have been another reasonable choice (however, with a smaller difference in SC-LC sensitivity to social context). Next, we explore SC sizes to identify an SC+LC sweet-spot to maximize the detection of EEG variance correlated with social context.

### 3.1 Isolating the contribution of social information to lexical prediction in LLMs

We explored the main parameter space of SC size (**Figure 3**) to identify context lengths that maximize the contribution of social speaker-related information when generating next-word predictions.

SC and LC classification accuracies were compared using FDR-corrected two-sided two-proportion z-tests. Classification performance increased as a function of context length. The LLM classification analysis indicated statistically significant differences in how much social information Mistral LLM can infer from SC (all sizes) and LC-4, with SC leading to a larger sensitivity to speech register and role across most SC sizes (**Figure 3A,B**).

For speaker role (podcast host vs. guest), accuracy was also consistently above chance for all context lengths excluding SC-1. SC-based predictions outperformed LC-4 predictions (50.65%) at context lengths of: 2 (52.12%), 4 (61.58%), 8 (63.61%), 16 (62.65%), and 32 words (69.67%) (**Figure 3A**). For speaker register (adult-directed vs. child-directed speech), Mistral achieved classification accuracy above chance level across all context sizes. SC-based accuracy exceeded LC-4 performance (62.69%) at window sizes of 2 (66.21%), 4 (70.53%), and 8 (71.22%) words (**Figure 3B**), but declined relative to LCs at larger windows, where the social information becomes absorbed in the LC (16-64 words; **Figure S2B**).

Further analyses were carried out to measure the influence of SC on the word surprisal and entropy. That influence was quantified by comparing time-series built with match and mismatch SC using the Pearson’s correlation score (**Figure 3C**). Lower correlation values, indicating a stronger influence of SC, were measured for entropy than for surprisal across all SC sizes. This difference was significant at every SC size (two-sided paired *t*-tests on Fisher z-transformed trial correlations, FDR-corrected; all *p* < 0.001). As this points at word entropy as the optimal candidate feature for measuring the EEG effect of SC, our main analyses of interest in this study focus on entropy. Further results on surprise can be found in **Figure S3.**

### 3.2 Isolating the influence of speaker context on the neural processing of speech at the word-level

Multivariate TRF models were fit, including acoustic (envelope, envelope derivative) and word-level features (word onset, word surprisal, word entropy). Word expectations values were quantified using Mistral LLM and the proposed SC+LM procedure using either matching or mismatching SC (**Figure 2**). Different SC sizes were explored (i.e., 1 word, 2 words, 4 words, 8 words, 16 words, and 32 words) to maximize sensitivity to social information and to validate the procedure. Specifically, we expected a small SC to bring a small or no contribution than a larger SC, with the caveat that an SC that is much larger than LC might reduce the relevance of the prediction to the local context, reducing both the prediction quality and the SC gain. We also expected effects small in magnitude, considering that word expectation in general bring small contributions to the explained EEG variance (Chalehchaleh et al., 2025). Nonetheless, these small gains were expected to be very robust across participants, given the paired within-participant nature of the SC match vs. mismatch contrast studied here^4^.

**Figure 2.**
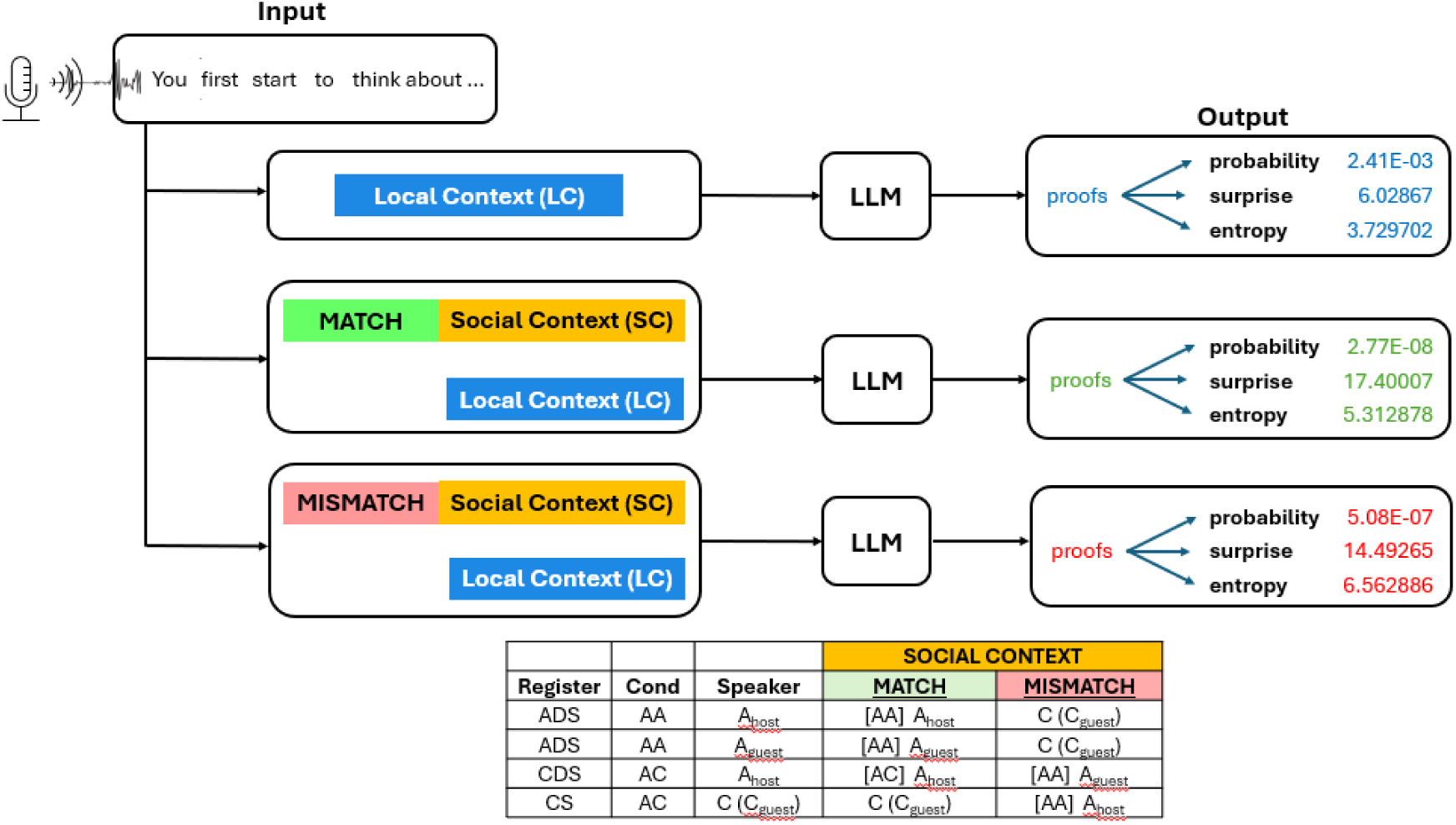
Modeling lexical predictions using different types of contexts changes LLM outputs. The podcast dialogue transcriptions that were fed into the LLM were modeled in 3 different ways to generate different values of lexical expectations (surprise and entropy): **(top)** using only the previous words before the predicted word as “local context” (Ip et al., 2025); **(middle)** using ‘matching’ (congruent) social context in addition to the local context; **(bottom)** using ‘mismatching’ (incongruent) social context in addition to the local context. The bottom table describes how ‘social context’ is defined and created in terms of speaker register per each experimental condition (adult-adult – AA or adult-child – AC conversations), speaker type (host or guest), and the congruent/incongruent social context (match or mismatch).

The results that follow measure the EEG prediction gain when fitting TRFs with acoustic features (envelope, envelope derivative), word onset, and word entropy (**Figure 4A**). EEG prediction correlations (average across all channels) when including word entropy resulted in a statistically significant main effect of SC size (repeated measures ANOVA; *F*(5,95) = 32.19, *p* < 0.001, Greenhouse-Geisser corrected, η² = 0.63) and congruency condition (SC_match_ vs SC_mismatch_) (repeated measures ANOVA; *F*(1,19) = 76.46, *p* < 0.001, Greenhouse-Geisser corrected, η² = 0.80). We also found a statistically significant interaction of SC size and congruency condition (two-way repeated measures ANOVA: *F*(5,95) = 18.26, *p* < 0.001, Greenhouse-Geisser corrected, η² = 0.49). A consistent EEG prediction gain of SC_match_ models emerged over their the SC_mismatch_ counterparts across all SC sizes, except size 16 (FDR-corrected Wilcoxon paired sign-ranked test, *p_SC=1_* = 0.002, *p_SC=2_* = 0.002, *p_SC=4_* < 0.001, *p_SC=8_* = 0.002, *p_SC=16_* = 0.41, *p_SC=32_* < 0.001). Single subject prediction correlations for SC-4, the highest SC_match_ vs SC_mismatch_ contrast, are visualized (**Figure 4B**) with the group means (SC_match_: 0.0891, SC_mismatch_: 0.0868). EEG prediction gains of SC_match_ over SC_mismatch_ were also detected at the level of single EEG-channel (**Figure 4C**). A one-sided, paired cluster-based permutation test comparing match and mismatch prediction correlations detected statistically significant prediction gains (*p =* ≤ 0.003) at four of the six SC sizes tested (SC = 2, 4, 8, 32). Statistically significant scalp areas spanned frontal, central, and centro-parietal regions.

**Figure 3.**
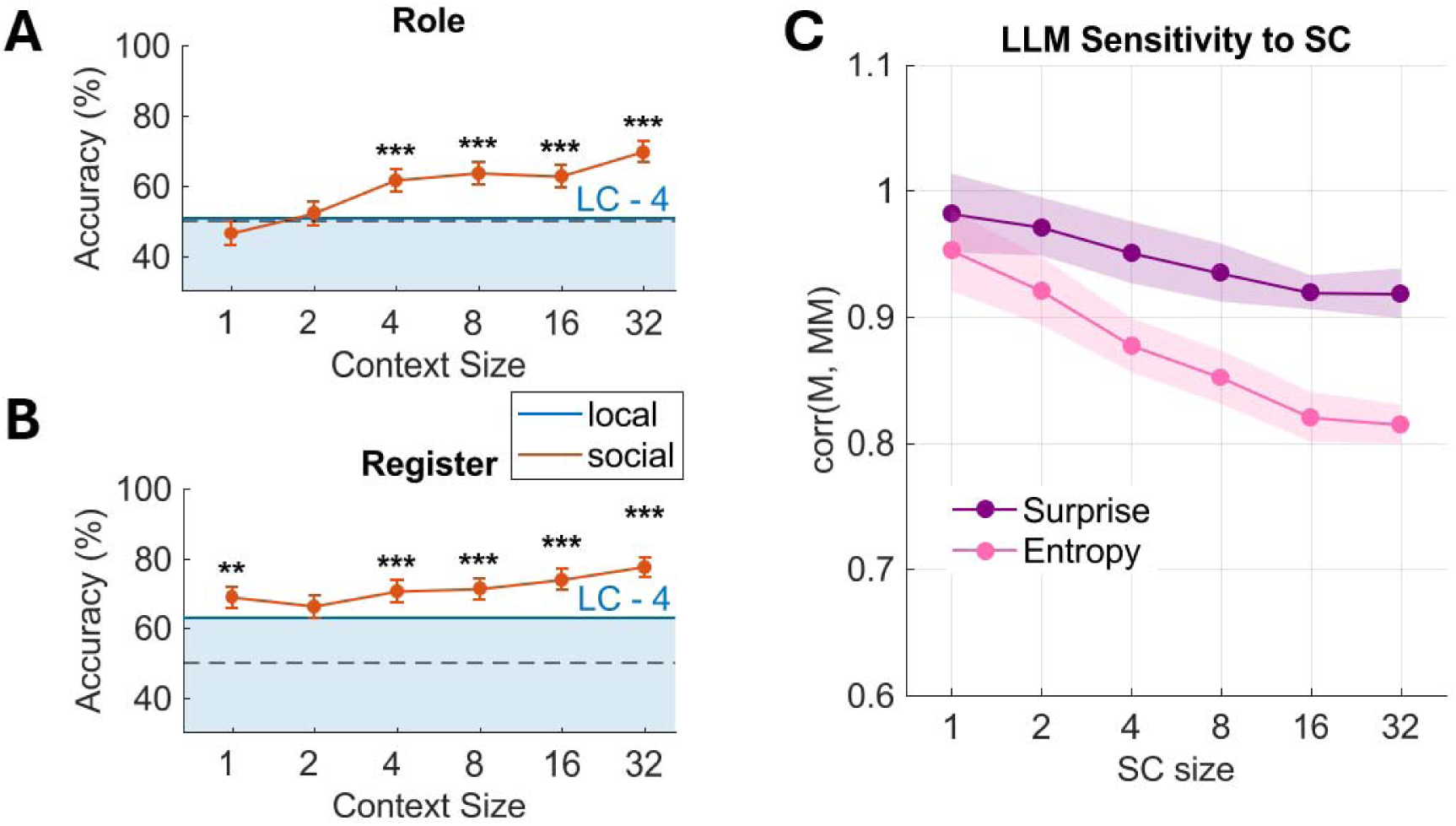
Selecting speaker-context parameters for isolating speaker-related contributions to lexical prediction using the LLM Mistral. Mistral was prompted to infer register and role information from a randomly generated set of 10,000 social and local contexts. Mistral’s accuracy was evaluated using two-proportion z-tests against ground truth labels for **(A)** speaker role (podcast host vs. guest) and **(B)** speaker register (adult-directed/adult speech or child-directed/child speech). The accuracy ceiling reached by “LC-4” is drawn as a solid blue line. **(C)** Raw average word surprisal (WS) and word entropy (WE) values for models conditioned on matching (congruent) versus mismatching (incongruent) social contexts with a fixed local context (LC) of 4 words were averaged across trials and then correlated across each SC size. Increasing the SC size, which introduces additional social context information, increases the contrast between congruent and incongruent models’ WS and WE measures.

**Figure 4.**
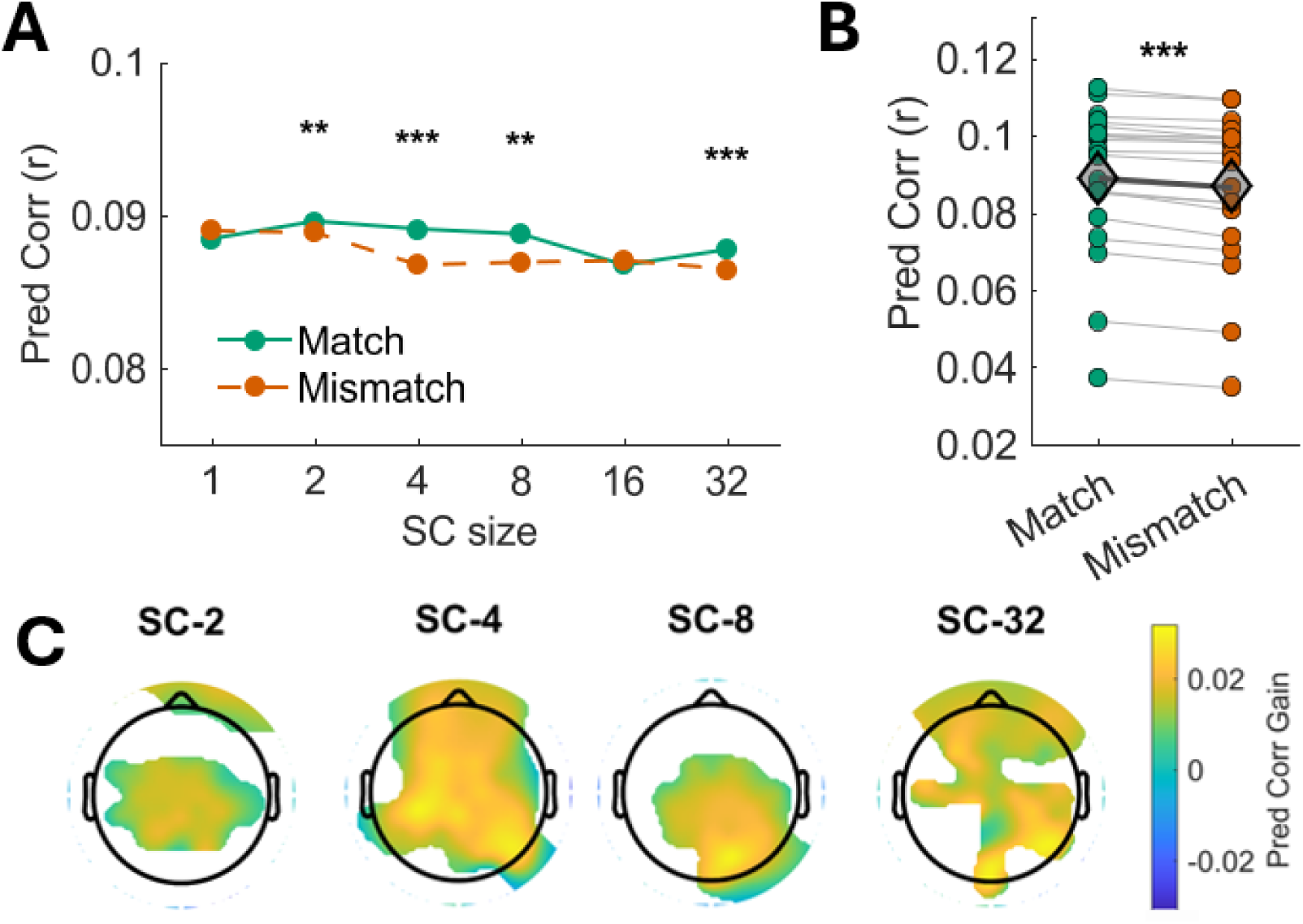
The brain uses a balance of speaker-specific and local contextual information to shape its linguistic predictions. Multivariate TRF models were fit on the acoustic features (envelope, envelope derivative, and word onset) and lexical expectation features, namely word surprisal (WS) and word entropy (WE), that were measured with congruent or incongruent speaker contexts. **(A)** The prediction correlations for models fit on Acoustics + (congruent and incongruent) WE are plotted, indicating the prediction gains on WE with social context information. **(B)** Single subject prediction correlations are plotted for SC-4 between the match (congruent) and mismatch (incongruent) models, with the group mean plotted using a gray diamond. **(C)** A cluster-based permutation test reveals those groups of electrodes in which congruent social context information has significant encoding gains in EEG over incongruent social context for models fit on Acoustics + WE.

We also present results from the equivalent analysis when also including word surprisal (**Figure S4**). A statistically significant main effect of SC size was detected (repeated measures ANOVA; *F(*5,95) = 19.54, *p* < 0.001, Greenhouse-Geisser corrected, η² = 0.51), no statistically significant main effect of congruency condition was measured (*F*(1,19) = 1.14, *p* = 0.30, Greenhouse-Geisser corrected, η² = 0.056), while a statistically significant SC size × condition interaction emerged (*F*(5,95) = 5.98, *p* < 0.001 (Greenhouse-Geisser corrected), η² = 0.24). Post-hoc one-sided Wilcoxon signed-rank tests (Match > Mismatch, paired, n = 20, FDR-corrected) showed significantly higher tracking for Match than Mismatch at SC only for SC sizes of 2 and 16 (both *p* = 0.002). Numerical results are also reported for LC with size 8, expecting weaker contrasts, as LC-8 captures more social information (**Figures S3B** and **S4**).

## 4. Discussion

This study presents a novel analytic procedure for measuring the influence of social information on the neural processing of speech at the word level. The framework combines LLMs, for formulating numerical hypothesis, and TRFs for hypothesis testing. Numerical results on a speech listening task involving podcast recordings demonstrate that small yet robust effects of social information on the EEG signal can be detected. This approach complements the neurophysiology toolkit for investigating the neurophysiology of speech, adding a procedure for probing neural correlates of word prediction that are influenced by social information.

Measuring the neurophysiology of speech with tasks approximating real-world experience has been intensively sought in recent years (Di Liberto & Ip, 2026). Advances in EEG technology and in analytic methods for recording EEG “in the wild” (Bleichner & Debener, 2017; Debener et al., 2015) or during socially interactive tasks (Dikker et al., 2017; Redcay & Schilbach, 2019; Zion Golumbic et al., 2013) give us new opportunities for probing speech communication in socially-relevant scenarios. Nonetheless, many of the analytic measurements that are available have been originally developed for controlled non-social scenarios. The present study contributes with a methodology that is thought specifically for studying scenarios with a diverse set of social contexts, offering an analytic approach for isolating the influence of social information within each condition and participant. The approach is conceptually different from the available analytic approaches, which would rely on comparing neural activity between conditions (as in **Figure S1**). The SC+LC procedure enables the formulation of alternative hypotheses on the neural reaction to words. This is similar to priming experiments, where the interpretation of ambiguous words or sentences can be altered by providing some information prior to the stimuli (Tikochinski et al., 2023; Yeshurun et al., 2017). One crucial distinction is that such priming experiments require speech material whose interpretation could substantially change due to prior information, and that this phenomenon was explored by using factual contextual knowledge as the prime. In contrast to that method, which requires a delicate design of the lexical content and speech material, our analysis focused on stylistic elements of the speech that change across register and conversational role. One point of strength of our procedure is its flexibility as, in fact, it could also be extended to include prior knowledge of facts on the speaker. This would constitute part of the priors available in a predictive processing framework, influencing the neural processing of upcoming words. While this is our initial intuition, alternative explanations based on semantic integration, memory retrieval, or other mechanisms of language comprehension may also account for contextual effects on speech processing (Brown & Hagoort, 1993; Huettig, 2015; Lewis et al., 2006; Nieuwland et al., 2018). Nevertheless, however, predictive processing gives us a solid intuition and a useful starting point for developing this first analytical framework to explicitly tease apart the local context information from other types of socially-relevant contextual information.

The proposed procedure combines SC and LC in ways that enable the isolation of EEG variance that is influenced by social information (i.e., by contrasting SC_match_ and SC_mismatch_). We would like to highlight that this is the objective of the procedure, which may come at the cost of lowering the overall quality of the LLM word prediction. From an LLM standpoint, this is counterintuitive, as the LLM cost function is to minimize prediction error. Our objective is different here, as LLMs can infer social information (e.g., on register and conversational role) from sufficiently long LC. As such, LC must be reduced in size to prevent the LLM from detecting social information from it, enabling the meaningful inclusion of SC segments.

For the purposes of this investigation, our selection of *Mistral-Instruct-7B-v.01* lent itself to this sensitivity, as it was fine-tuned on conversational datasets. Our SC+LC procedure can be applied using other LLMs, potentially leading to different results, reflecting the particular sensitivities of the model. For example, an LLM may be more sensitive than another to particular aspects of speech register, such as lexical diversity and word frequency distributions. As such, it is crucial to estimate the particular sensitivities of the model for the selected SC and LC sizes, for a given LLM and text material. That could be done via a zero-shot classification, as in **Figure 3**, but other methods could also be considered, such as studying how contextual information influences the contextual word embeddings inside the model. In either case, please note that LLMs are taken as a convenient approach for generating statistically-relevant numerical hypotheses, and are not intended as an accurate model of brain functions (Caucheteux & King, 2022; Mischler et al., 2024).

Multivariate TRF models on the feature set combining acoustics and WE revealed that conditioning lexical expectations on congruent social context significantly and consistently outperformed those based on incongruent social information. This result is in line with our initial motivation, where we postulated that our brains combine local and social information when building next-word predictions, where social information here included speaker register and conversational role. Statistical analyses were carried out on the prediction gains after averaging all EEG channels and are, as a consequence, small values. Prediction gains for single channels, instead, reach values of 0.02 and above (average across participants), which is a relatively large correlation gain that is comparable in magnitude and topographical locations with word-level EEG prediction gains in other datasets (for example, see Figure 3F in (Chalehchaleh et al., 2025)). More work on other datasets is necessary to determine the effectiveness and generalizability of this approach at detecting different types of social information, tasks, and individual socio-cognitive skills.

Although the present framework is limited in how we define “social information,” in the context of communication and how much of social cognition it captures relative to what lies at the heart of communicative behaviors, it constitutes a step towards more “dialogue-aware” models of speech processing by incorporating information about interlocutors and speaking styles into computational estimates of lexical prediction. Broadly, these pieces of information are important and taken into account during the active coordination that interlocutors perform to create common ground, and the theory of mind principles that guide partner modeling and how interlocutors may transmit their communicative intentions. Other definitions of social context refer to the social framing of monologues or dialogues (Abassi & Zatorre, 2025), perceived live listening versus prerecorded listening tasks (Rice & Redcay, 2016), and which roles interlocutors play in a conversation (Bell, 1984). Our proposed framework contributes one possible way of operationalizing social context, namely speaker register and role, but we advocate for future research in this framework to capture even more dimensions of what can be considered “social context.” Certainly, other social context information used by listeners and interlocutors in a speech interaction could include power dynamics, communicative goals, common ground, familiarity with speakers, and can be modeled at different levels of the linguistic hierarchy in prosodic or paralinguistic features.

## Supplementary

**Figure S1.**
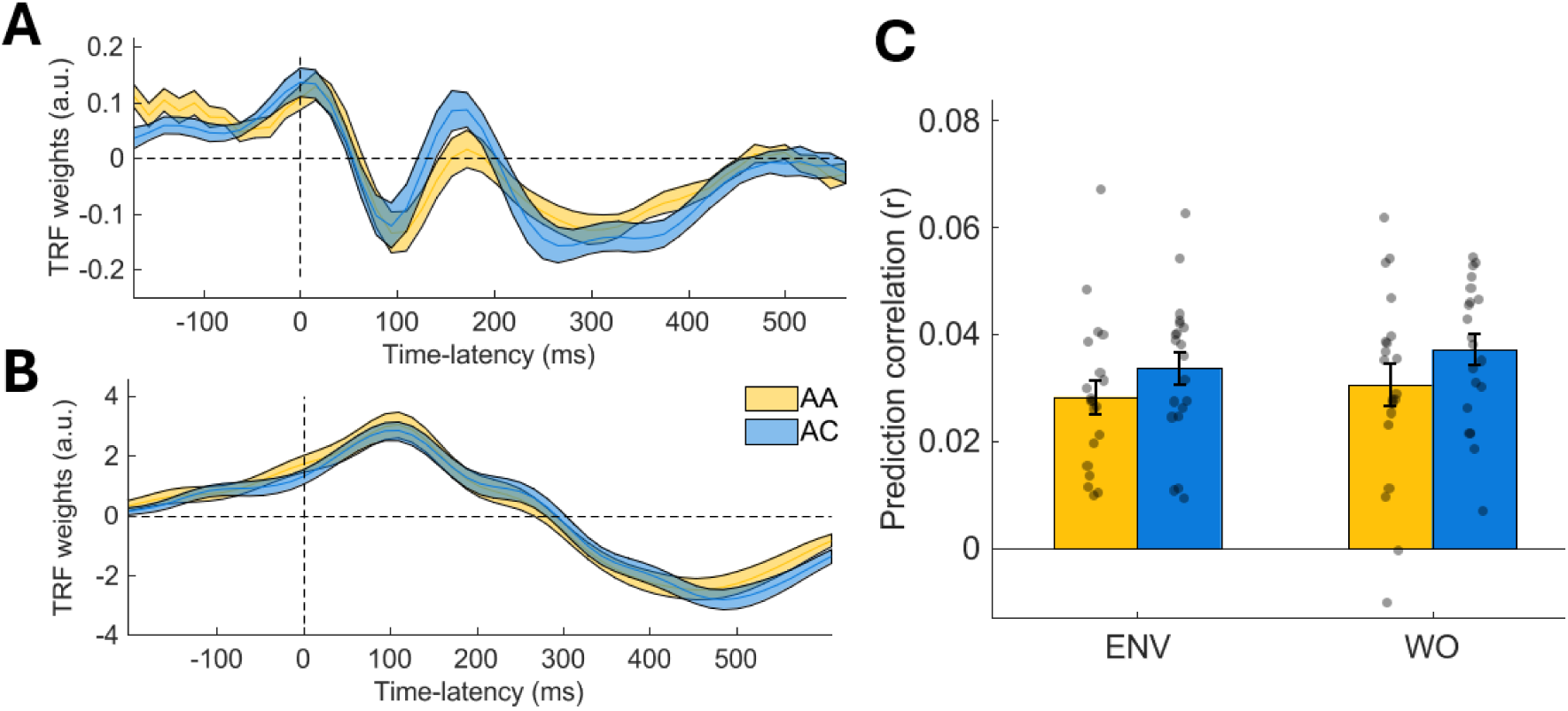
Weights of the envelope and word onset features, split across ADS and CDS/CS conditions. **(A)** Envelope weights **(B)** Word onset weights. **(C)** Prediction correlations.

**Figure S2.**
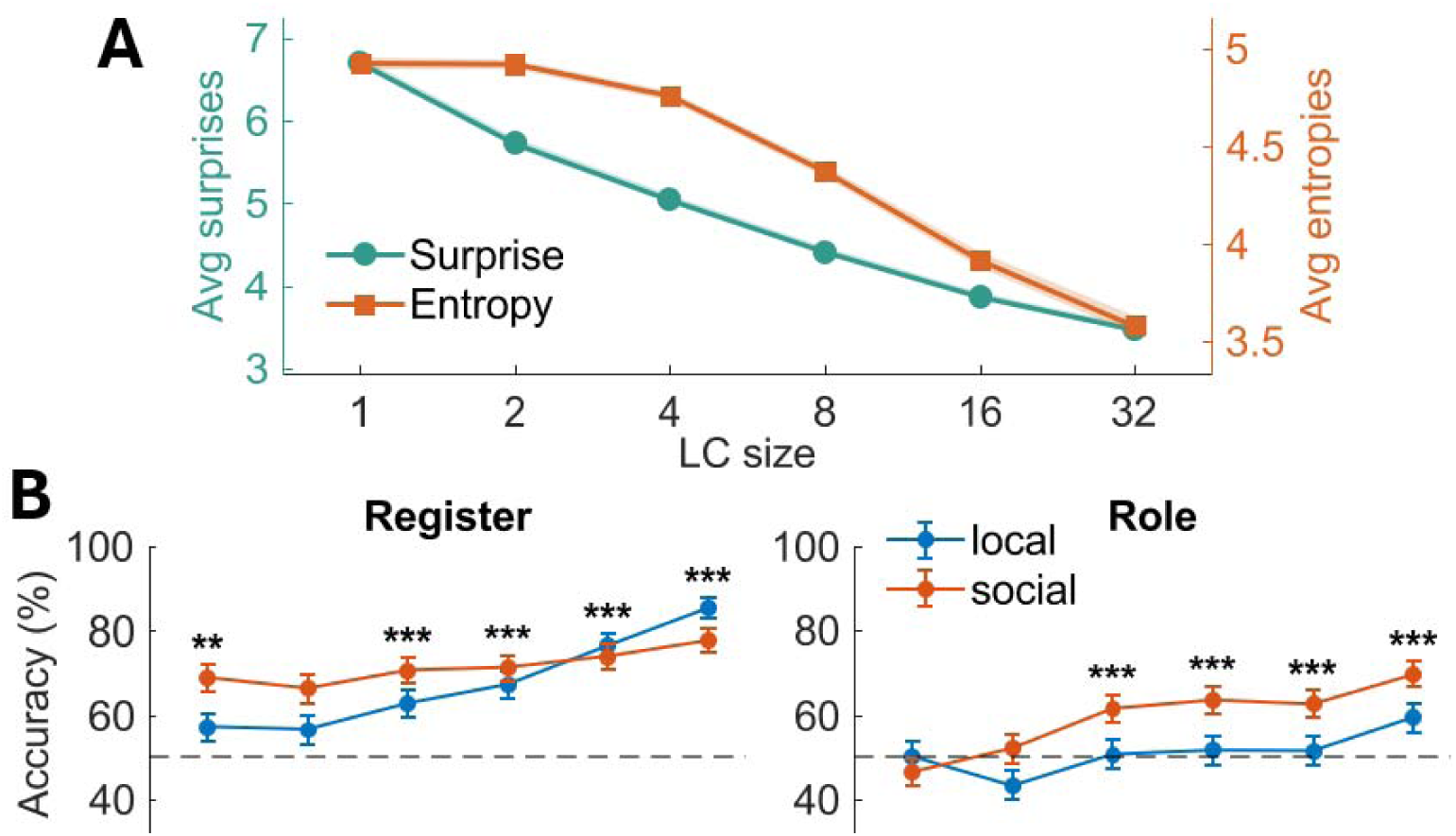
Social speaker information can be isolated from local context information using the LLM Mistral. **(A)** The average surprise and entropy values for different context lengths of local contexts are plotted. Mistral was prompted to infer register and role information from a randomly generated set of 10,000 social and local contexts. **(B)** Mistral’s accuracy was evaluated against ground truth labels for **(left)** speaker register (adult-directed/adult speech or child-directed/child speech) and **(right)** speaker role (podcast host vs. guest). This extends the data from Figure 3.

**Figure S3.**
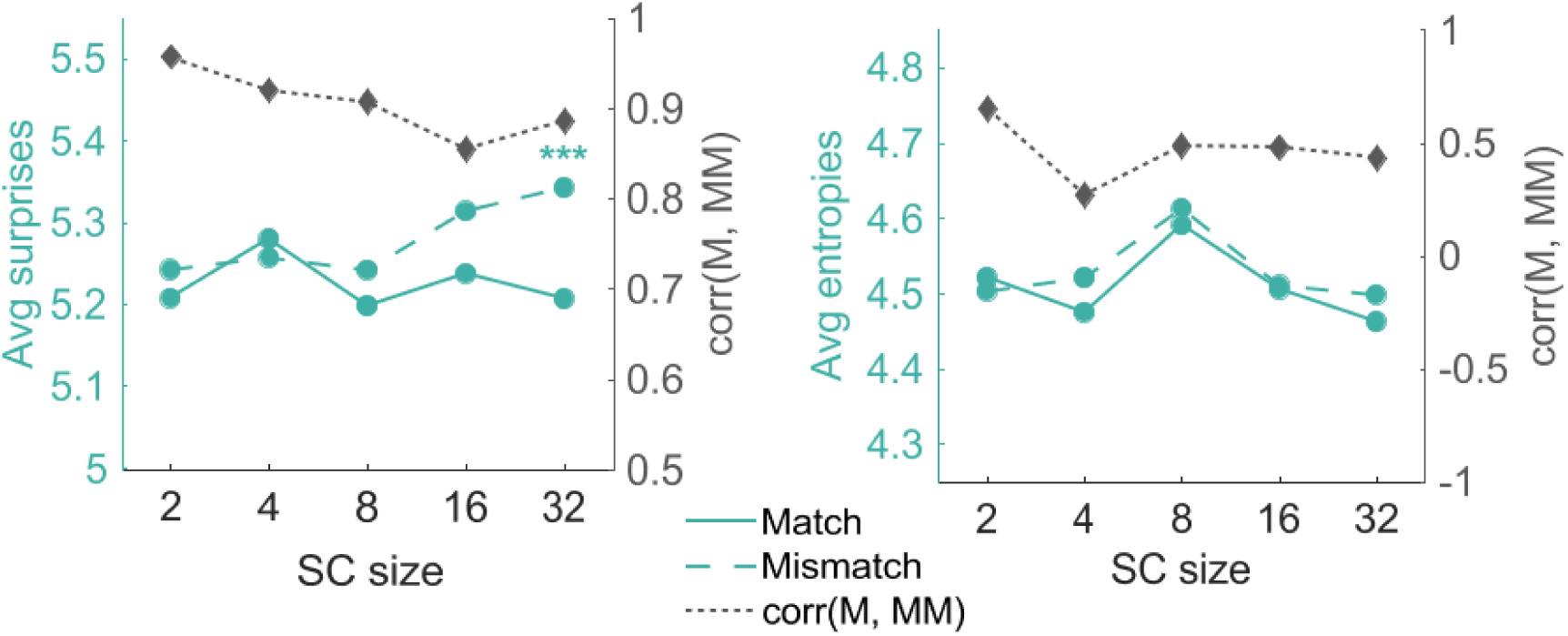
Raw average surprise values are shown for models conditioned on matching (congruent) versus mismatching (incongruent) social contexts with a fixed local context (LC) of 4 words. As SC size increases, introducing additional social context information, Mistral’s predictive performance declines, reflected in higher overall surprise (left, teal y-axis) and an increased divergence between congruent and incongruent models (right, gray y-axis).

**Figure S4.**
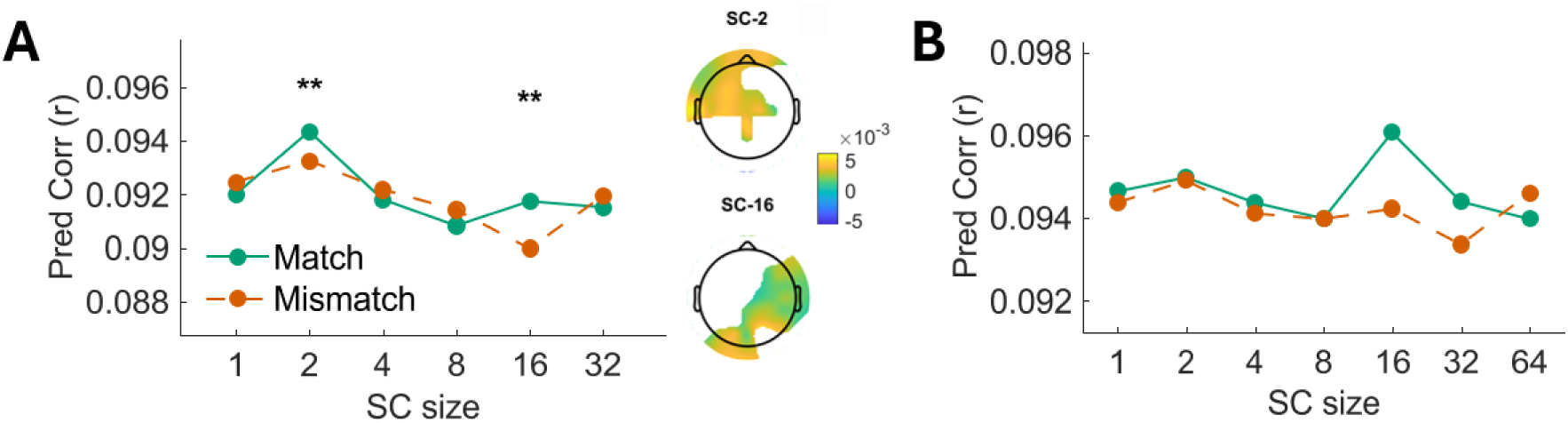
Acoustics + Surprise + Entropy mTRF prediction correlations for LC-4 and LC-8. **(A)** The mTRF feature set that included acoustic features combined with surprise and entropy revealed prediction gains of the congruent SC models over their incongruent SC counterparts for LC-4 at SC-2 and SC-16. The corresponding topographies reveal a significant cluster in the left fronto-temporal region for SC-2 and the right-lateralized centro-parietal region for SC-16. **(B)** No significant gains in this feature set emerged for LC-8. This pattern was highly consistent across SC sizes from 1 to 32, with the posterior-occipital cluster surviving correction at every level tested (all p ≤ .0002) in LC-4.

**Figure S5.**
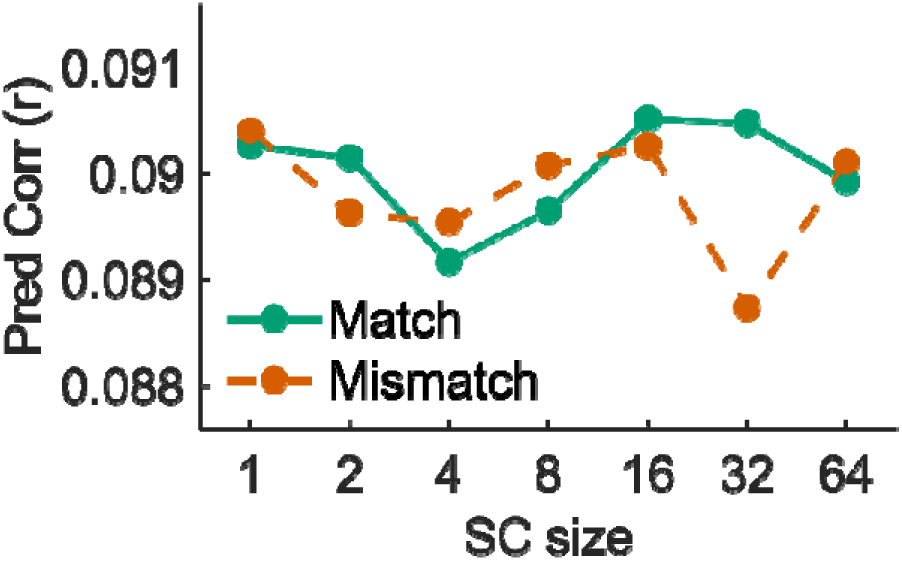
Acoustics + Entropy mTRF prediction correlations for LC-8. The mTRF feature set that included acoustic features and word entropy revealed no significant prediction gain of the congruent SC models over their incongruent SC counterparts.

## Footnotes

1 https://laist.com/podcasts/servant-of-pod/kids-podcasts-a-true-alternative-to-screen-time; https://creators.spotify.com/pod/profile/martine-severin/episodes/31--How-to-Become-an-Epic-Improv-Comic-with-Joy-Dolo-e1js9c2

2 https://github.com/CNSP-Workshop/CNSP-resources/tree/main/CNSP/exampleScripts

3 https://cnspworkshop.net/

4 similar to the widely studied selective attention decoding, where small differences in prediction correlations between attended and unattended streams can be extremely consistent and lead to strong decoding classification accuracies (O’Sullivan, J. A., Power, A. J., Mesgarani, N., Rajaram, S., Foxe, J. J., Shinn-Cunningham, B. G., Slaney, M., Shamma, S. A., & Lalor, E. C. (2015). Attentional Selection in a Cocktail Party Environment Can Be Decoded from Single-Trial EEG. *Cerebral Cortex (New York, N.Y.: 1991)*, *25*(7), 1697-1706. https://doi.org/10.1093/cercor/bht355

